# Biogeography and Microscale Diversity Shape the Biosynthetic Potential of Fungus-growing Ant-associated *Pseudonocardia*

**DOI:** 10.1101/545640

**Authors:** Bradon R. McDonald, Marc G. Chevrette, Jonathan L. Klassen, Heidi A. Horn, Eric J. Caldera, Evelyn Wendt-Pienkowski, Matias J. Cafaro, Antonio C. Ruzzini, Ethan B. Van Arnam, George M. Weinstock, Nicole M. Gerardo, Michael Poulsen, Garret Suen, Jon Clardy, Cameron R. Currie

## Abstract

The geographic and phylogenetic scale of ecologically relevant microbial diversity is still poorly understood. Using a model mutualism, fungus-growing ants and their defensive bacterial associate *Pseudonocardia*, we analyzed genetic diversity and biosynthetic potential in 46 strains isolated from ant colonies in a 20km transect near Barro Colorado Island in Panama. Despite an average pairwise core genome similarity of greater than 99%, population genomic analysis revealed several distinct bacterial populations matching ant host geographic distribution. We identified both genetic diversity signatures and divergent genes distinct to each lineage. We also identify natural product biosynthesis clusters specific to isolation locations. These geographic patterns were observable despite the populations living in close proximity to each other and provides evidence of ongoing genetic exchange. Our results add to the growing body of literature suggesting that variation in traits of interest can be found at extremely fine phylogenetic scales.

## Introduction

The microbial world encompasses a vast amount of phylogenetic, genomic, and ecological diversity (1). However, linking sequence-based metrics of diversity to differences in ecological characteristics remains difficult (2). One major challenge is that a given taxonomic level is generally defined much more broadly in microbes than in eukaryotes. For example, even “closely-related” bacterial groups such as *Salmonella* and *E. coli* are estimated to have diverged approximately 100 million years ago (3). In the genus *Streptomyces*, studies of strains with nearly identical 16S rRNA gene sequences display both diverse antibiotic resistance and resource use phenotypes (4), and retain spatial distributions influenced by glacial movement in the last ice age (5). Additionally, predicting and investigating ecologically relevant phenotypes in microbes is extremely challenging. For example, *Vibrio cyclotrophicus* strains isolated from different size oceanic organic particles displayed divergence in only a small number of specific genes (6). Even with these data as a guide, significant experimental examination was required to reveal subtle but ecologically significant phenotypic differences that helped explain their physical distribution (7).

Although the environmental forces driving the distribution of fine-scale microbial diversity are poorly understood for most taxa, those that are associated with extreme environmental conditions (8) or with eukaryotic hosts (9) can be used to address biogeographical and population-scale ecological questions more readily. Lineages of bacteria from the actinobacterial genus *Pseudonocardia* that form a defensive mutualism with many fungus-growing ant species (10, 11) provide a useful model system to do this. These filamentous spore forming bacteria grow on the external surface of the ants, where they produce natural products that inhibit the growth of *Escovopsis*, a co-evolved pathogen of the ant’s fungus garden (12). Transmission occurs from one ant to another within the first hours of adult life (13). Queens generally carry the bacteria with them when forming a new colony, but there is phylogenetic evidence of host switches over evolutionary time and potential acquisitions of new symbionts from the environment (14). These bacteria have also become a source of novel small molecules (15, 16).

We hypothesized that the genomic diversity of ant-associated *Pseudonocardia* and their natural product repertoire would vary with host biogeography even over relatively small distances. We investigated this using genomes of *Pseudonocardia* isolated from *Apterostigma* ants within a 20 km transect on and around Barro Colorado Island (BCI) in Panama which formed a single clade in a recent multilocus phylogeny (17). This island was formed from a hilltop that became isolated due to the influx of water that created the Panama Canal. Using a combination of comparative genomic and population genetics, we investigated micro-scale diversity of genome content, genetic exchange, and metabolic potential across kilometer-scale geographic space.

## Results

The 46 *Pseudonocardia* genomes had a median size of 6.68mb, with a median GC content of 73.42% and 7,253 open reading frames (ORFs) and inferred RNA coding regions. The number of ORFs is likely inflated by the poor assembly quality of illumina sequenced genomes, as the median number of coding regions in high quality genomes (fewer than 10 contigs) is 5935 (Supp Table 1). These genomes had extremely high sequence similarity, with on average 99.4% nucleotide identity in core genes (Figure 1). We identified 280,007 dimorphic SNP positions by mapping all genomes in the clade to the complete genome of *Pseudonocardia* sp. EC080625-04, also associated with *Apterostigma*. Putative population assignments using the SNP data matched the isolation locations for most strains, with two groups of BCI isolated strains, BCI-A (light blue) and BCI-B (dark blue), and a number of strains from the mainland around Pipeline Road (PLR), labeled PLR-A (green), PLR-B1 (red), and PLR-B2 (orange) (Figure 1a, b).

**Figure 1:**
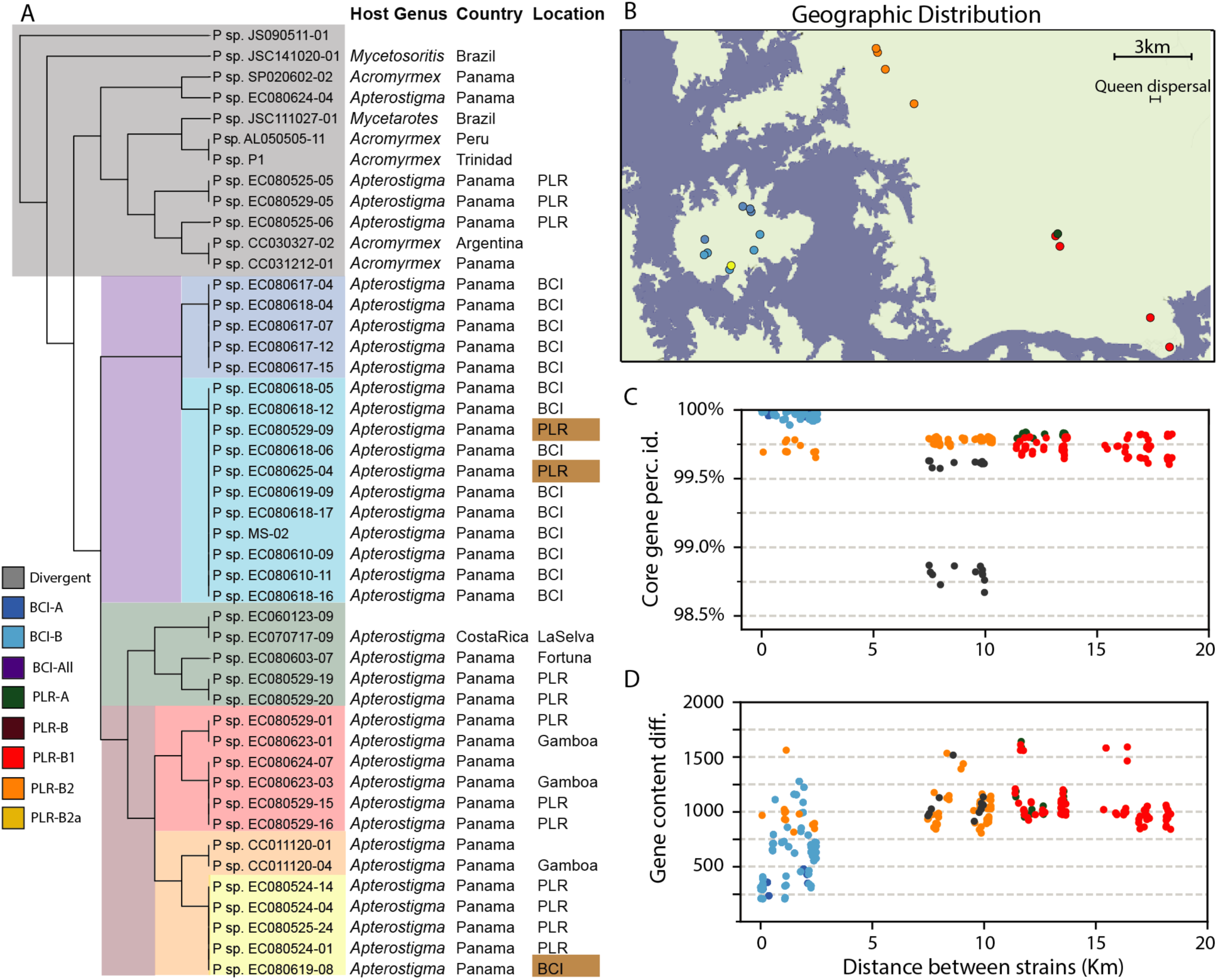
SNP-based clustering of ant-associated *Pseudonocardia*. A. fineSTRUCTURE clustering of *Pseudonocardia* strains using 270,000 dimorphic SNP positions. Ant host and isolation location are shown to the right. Lineage colors are used throughout the study. Strains whose local isolation location conflicts with others in their cluster are label in tan. B. Isolation locations based on available GPS data are shown for *Pseudonocardia* strains in the area around Barro Colorado Island. Strains are colored by fineSTRUCTURE population. C. Core gene percent identity between BCI cluster strains (purple lineage in A) and all strains with GPS coordinates. Points are colored by strain population. D. Raw number of gene content differences between BCI cluster strains and all others with GPS coordinates.

The SNP-based clusters of BCI strains largely matched their geographic distribution on the island, with the less diverse BCI-A subpopulation occupying the northern and western part of the island. PLR strains also clustered by geographic distribution, with PLR-B2 being in the northern part of our sampling area, while the sympatric populations PLR-A and B1 were isolated 7.6 km further south. Finally, three strains of PLR *Pseudonocardia* grouped with *Pseudonocardia* isolated from other fungus-growing ant species or from *A. dentigerum* colonies from other countries (grey). Sampling of these lineages is more limited in our dataset, with many strain clusters consisting of only one or two genomes. There were three strains whose cluster assignment conflicted with their isolation location: two mainland-isolated strains clustered in BCI-B (*Pseudonocardia* sp. EC080625-04 and EC080529-09) and a single BCI-isolated strain (EC080619-08) clustered with PLR-B2 (Figure 1a).

The number of polymorphic sites between genomes in reference-aligned regions ranged from 300 to more than 200,000 (Supp Fig 1). Overall, core gene percent identity between isolates from the BCI populations and mainland genomes matched their geographic locations (Figure 1c, Supp Fig 2a). Except for the isolate that grouped with the mainland strains, BCI strains had very high sequence similarity with other isolates from the island, and lower core gene similarity to mainland strains. Nearly all strains from the area around BCI shared core gene percent identities above 99.5%, except for three strains with a percent identity to the BCI strains of 98.8%.

Genome content diversity also followed a geographical pattern, with BCI strains generally being similar in genome content (Figure 1d, Supp Fig 2b). Gene content of strains from the BCI populations differed from mainland strains by around 15%. Between BCI strains, gene content differences range from 188 to 2,063 genes. The pan-genome size of these closely-related *Pseudonocardia* strains was relatively small, with 6,617 gene families present in at least one genome (Supp Fig 3). The core genome is approximately 3,200 gene families, which represents about half of the total genes found in most *Pseudonocardia* genomes in our dataset.

Analysis of contigs mapping to the reference genome provided insight into the source of gene content variation between these genomes. The vast majority of contigs either mapped to the reference across nearly their entire length, or failed to map almost entirely. Cases of only part of a contig mapping to the reference were very low, at only 3.5% of contigs with more than 10% and less than 90% mapping to the reference. Further, non-mapping contigs had a lower GC content than mapping contigs, at 72% versus 74% respectively (t-test statistic −19.7, p-value 2.82E-85). Many pathways associated with secondary metabolism were over-represented among genes that did not map to the reference genome, along with genes for the degradation of xenobiotics and a number of amino acids, and transposon/phage genes. Genes involved in many core biological functions and metabolic pathways were under-represented among genes that did not map to the reference, as expected for conserved functions.

Since the production of antimicrobial compounds is thought to be the primary ecological role of ant mutualist *Pseudonocardia*, we investigated the diversity and distribution of natural product biosynthetic gene cluster families (BGCs) among the sampled *Pseudonocardia* populations (Fig 2). We identified 27 BCG families, 7 of which were only common in the southern BCI population BCI-B. Four additional BGCs were significantly enriched among BCI populations, including a siderophore and two type-II polyketide BCGs. Mainland-specific BGCs included a predicted lassopeptide found only in isolates from the southern part of the mainland area, and a nonribosomal peptide synthetase found only in the northern mainland area. Multiple BGCs, including an ectoine, an oligosaccharide, a terpene, and a nonribosomal peptide synthetase were present within mostly distant populations while being underrepresented or absent in the PLR and BCI populations.

**Figure 2:**
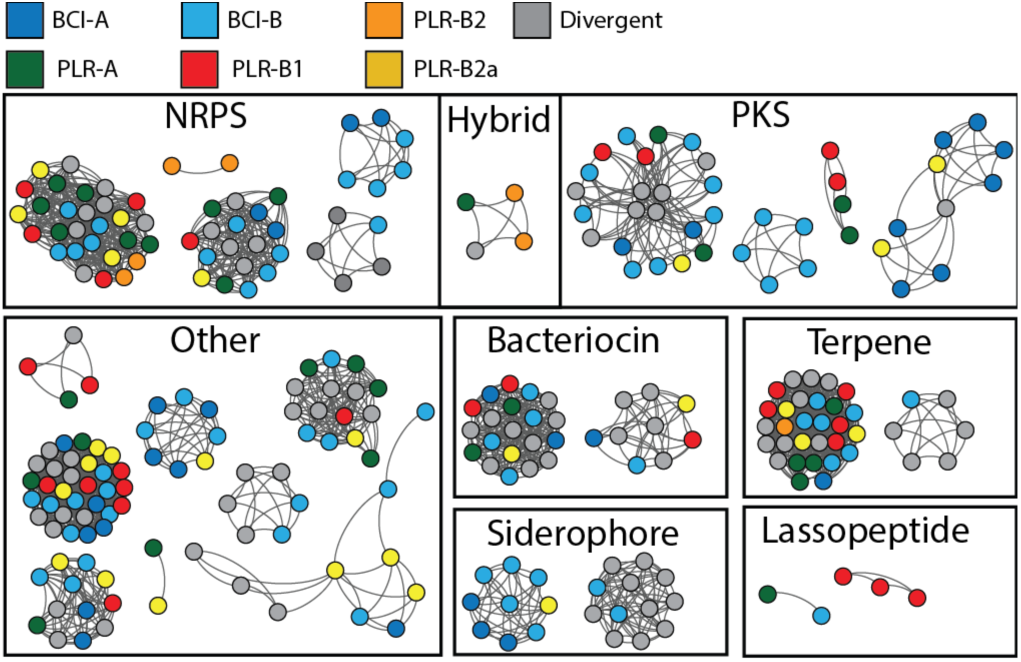
Recurrent natural product biosynthetic gene clusters. Each node represents a contiguous set of genes that are part of a natural product biosynthesis cluster. Edges represent at least 80% nucleotide sequence identity and 50% coverage between gene sets and are unweighted. Nodes are colored by their population of origin (see Figure 1).

Recombination events were non-randomly distributed across the chromosome, with some regions unaffected by recombination events and others affected by as many as 50 events. Nearly all large blocks of high recombination density overlap with transposons and hypothetical proteins. The amount of recombination detected in each strain varied considerably, from 0.1% to 14.1% with a median of 5.9% (Figure 3A). Genes that formed monophyletic clades within each genome cluster, and therefore were not significantly affected by recombination between populations, were generally distributed across the chromosome in BCI populations (Figure 3b). However, in the PLR lineages there was a pronounced bias. The 248 and 784 monophyletic genes in the red and orange lineages, respectively, were found almost exclusively in the middle portion of the reference genome chromosome, surrounding the origin of replication. This bias was reversed among the 581 monophyletic genes in PLR-B2a (gold). In this lineage, the outer portions of the chromosome were enriched in monophyletic genes.

**Figure 3:**
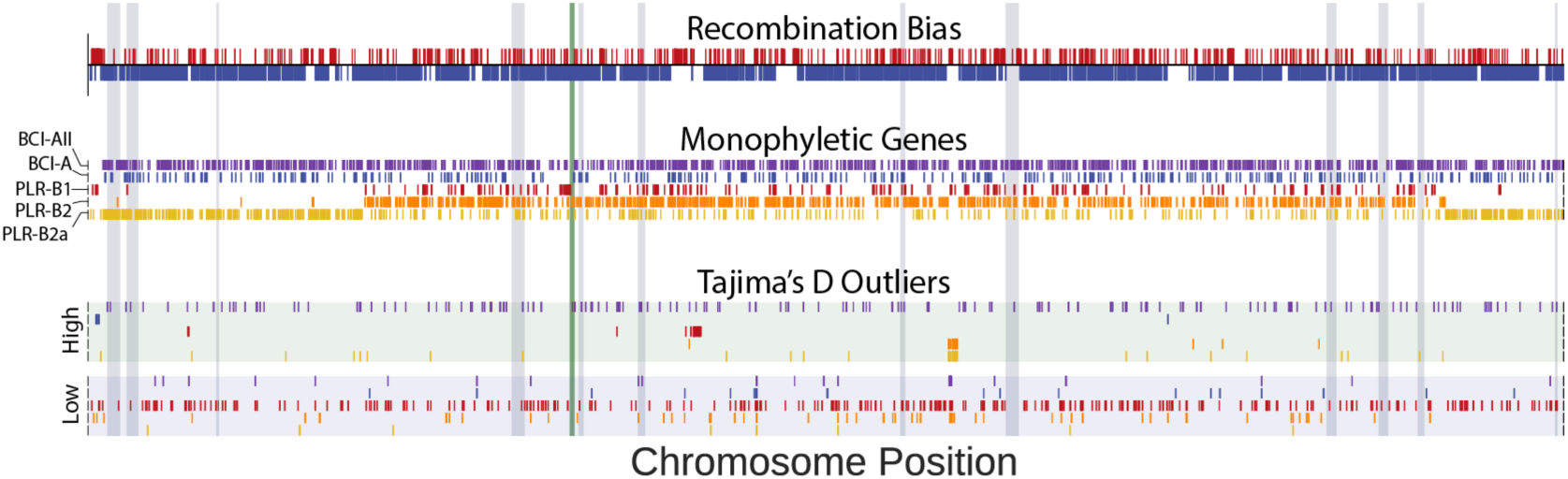
Chromosome map of recombination events and divergent genes. All genomic contigs were mapped to the complete genome of *Pseudonocardia* EC080625-04. Grey regions indicate natural product biosynthesis clusters. The vertical green bar indicates the inferred origin of replication. Populations are colored as in Figure 1. Recombination track shows chromosome regions enriched (red) or depleted (blue) of recombination events. Monophyletic Genes track shows the chromosomal location of genes whose phylogenies form monophyletic groups for each population. Tajima’s D Outliers track shows the location of genes whose Tajima’s D values deviate by more than 2 standard deviations from the mean of each population.

The ancestral node shared by BCI-A and B (purple) had the highest number of monophyletic genes, at 1,023, while BCI-A and BCI-B were distinguished by 344 and 43 genes, respectively (Figure 4). Functional categories enriched among BCI monophyletic gene trees include xenobiotics degradation (odds ratio 1.69), tryptophan metabolism (odds ratio 2.18), ABC transporters (odds ratio 1.67), and nucleotide excision repair (odds ratio 4.64). KEGG gene categories enriched among the monophyletic genes in PLR-B2 (red) include secondary metabolism (odds ratio 1.82) and aminobenzoate degradation (odds ratio 2.21).

**Figure 4:**
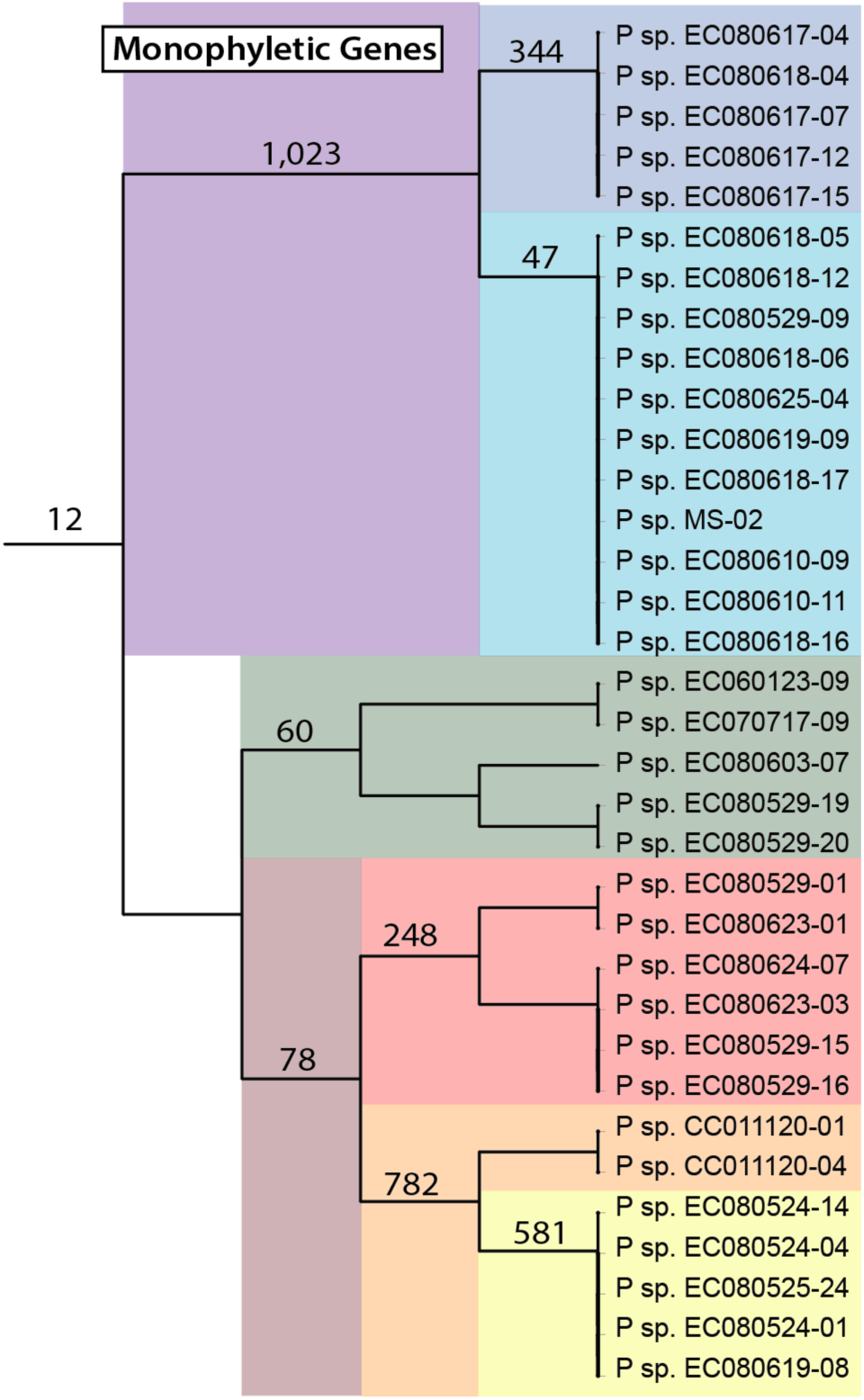
Pipeline road region population-specific gene divergence. The number of genes that exhibit monophyly for the strains in each lineage containing at least 5 strains are shown on the appropriate branch.

We calculated Tajima’s D (18) values (TD) for all genes with greater than 1% polymorphic sites in various strain clusters (Figure 5) to investigate gene-level sequence diversity. The full population-scale dataset had a mean TD value of −0.444 across 4,225 genes. The BCI strain groups had much lower genetic diversity, with 404 genes containing enough polymorphic sites to calculate TD and a mean value of −1.41. While the PLR-A cluster also had a low TD value, PLR-B (dark red) had a median TD value of −0.03, and the PLR-B1, and PLR-B2 had mean TD values of 1.28 and 0.28, respectively.

**Figure 5:**
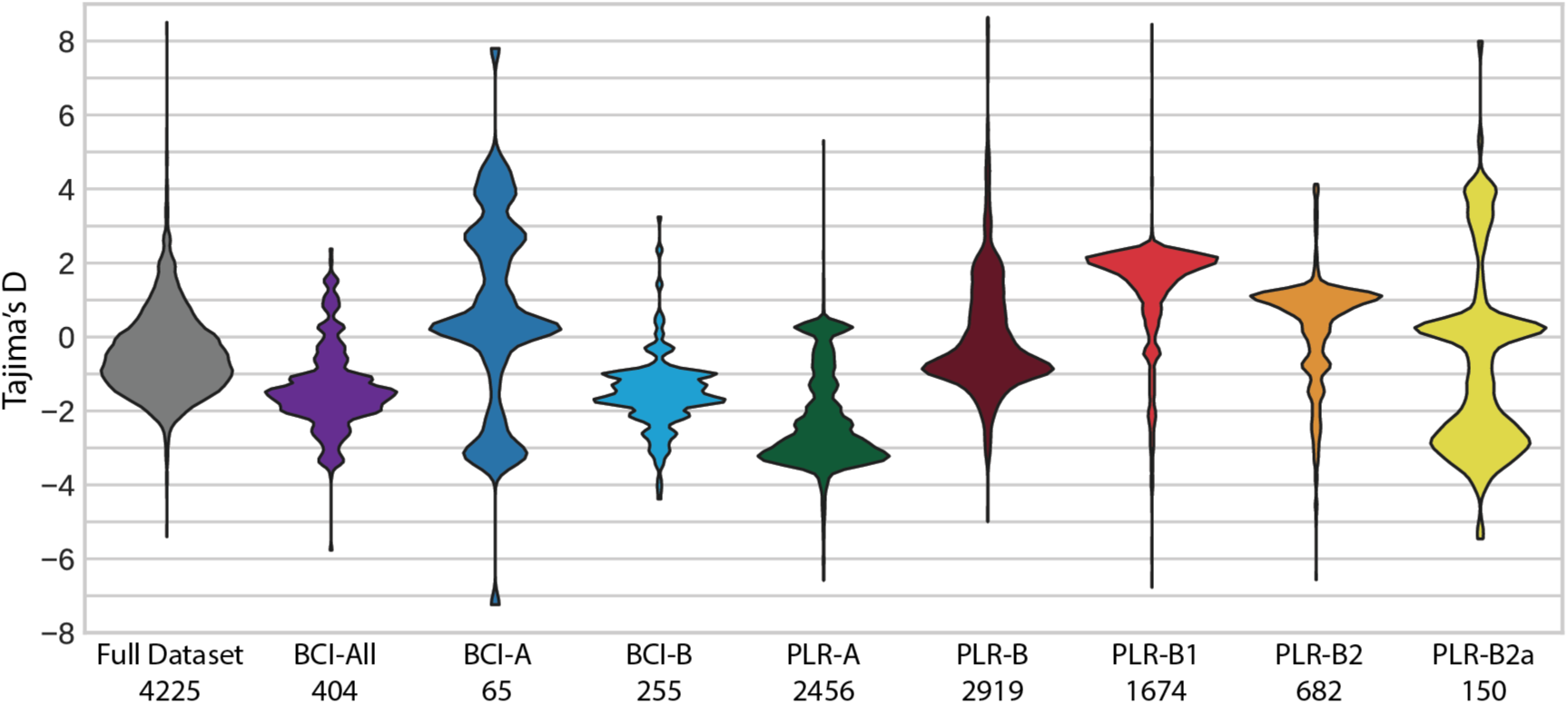
Genetic diversity of conserved genes. Violin plot of the distribution of Tajima’s D values for the specified strain group in genes containing at least 1% polymorphic sites. The number below each strain group name indicates the number of genes analyzed.

We also investigated genes with TD values that deviate from the mean by more than 2 standard deviations in each population, and in all genomes combined. In most individual populations these genes were spread across the genome, except for those in PLR-B (dark red) that cluster near the origin, and a cluster of genes with high TD in PLR-B2 (orange) and B2A (gold) that appear to be part of a mobile element (Figure 3). Across each individual population, genes containing the PFAM (19) domain of unknown function DUF222 were significantly enriched among genes with unusually low TD values (Fisher’s exact test, odds ratio 50.02, p value 1.2E-9). Eleven genes in EC080625-04 contain this domain, eight of which are in the lowest 5% of TD values in at least one population and seven of which are in the lowest 5% in at least three populations. Two other PFAM domains were enriched in the low TD genes, one of which is transposase related (DDE_Tnp) and another a repeat domain of unknown function (RCC1_2).

When the full set of genomes were analyzed as a single dataset, a number of gene clusters showed high TD values. These included several transporters and signaling proteins, along with a cluster of genes involved in exopolysaccharide and cell envelope biosynthesis. Genes with low TD values were more scattered and included a putative prophage and several transposases in addition to the aforementioned DUF222 domains. There were several clusters of low TD genes, including a number of genes in a type-VI secretion system. A number of core genes also show abnormally low TD, including FtsQ and cytochrome C oxidase subunit I. When the two BCI populations were analyzed together, low TD genes included a large number of hypothetical proteins along with two secretion-system associated proteins, one type IV VirD4 family and one type VII EccB family. High TD genes included an MT0933-like antitoxin protein along with a range of hypothetical proteins.

## Discussion

Population genomic analysis of *Pseudonocardia* associated with fungus-growing ants provides a significant contribution to the growing body of literature demonstrating that extremely closely-related bacteria, well within standard OTU definitions, can exhibit both gene content diversity and distinct signatures of population dynamics (4–7). Geography plays a clear role in structuring the genome sequence similarity and gene content diversity of *Pseudonocardia*, with BCI isolates being highly similar and forming a single lineage that is distinct from most mainland strains. The only BCI-isolated strains that do not share very high sequence similarity to the others instead share high identity with some mainland isolates, suggesting either continuing migration of ant hosts between the island and mainland or sustained coexistence of two lineages since the formation of the island. Similarly, two mainland strains share high sequence similarity with the island strains.

The distribution of natural product BGC families also follows isolation location at this fine geographic scale, as a number of BGC families were found only in BCI strains. This may be due to geographic diversity of fungus-growing ant pathogens, as dynamic natural product potential is likely important for maintaining effective inhibition of pathogenic fungi. The acquisition of several of these clusters by the recent migrant strain *Pseudonocardia* sp. EC080619-08 may suggest either rapid acquisition of ecologically relevant genes after host migration, and/or strong selection from pathogens or other environmental conditions prevent the colonization of BCI by ant hosts whose *Pseudonocardia* lack the ability to produce particular small molecules. More samples from both mainland and BCI populations would help address this question by providing a more comprehensive view of BGC family conservation and diversity across locations. Studies on migration and survival of new *Apterostigma* ant colonies would also shed light on the dynamics of host dispersal and survival in new geographic areas.

The large number of monophyletic gene trees that separate the BCI lineage from the mainland lineage provide strong support for genetic isolation of many loci, despite ongoing recombination. This observation is particularly important when investigating evolutionary independence between closely related bacterial lineages. As homologous recombination occurs in relatively small, non-random stretches of DNA rather than uniformly across the entire chromosome, bacterial populations can be genetically isolated at some loci while recombining at others (6). This partial genetic isolation model is supported by the strongly biased chromosomal distribution of monophyletic genes in the PLR-B lineages, overlapping with the origin of replication. Such a pattern is consistent with divergence of the core genome despite continued genetic exchange in accessory genes primarily located on the chromosomal periphery, similar to physical distribution of accessory genes on the chromosomes of other Actinobacteria such as *Streptomyces* (20). Reversal of this trend in PLR-B2a may suggest recombination events or divergence of accessory genes unique to the PLR-B2a lineage. This creates an abundance of monophyletic genes in the periphery of the genome in PLR-B2a strains while they remain less distinguishable from other members of the PLR-B2 lineage in more conserved genes.

Our identification of diverse population dynamics and natural product biosynthetic potential, structured by their geography, suggest this lineage of closely related ant-associated *Pseudonocardia* contains multiple distinct populations. Categorizing these microbes by sequence similarity alone (21) would incorrectly infer that they are biologically equivalent. Thus, understanding the micro-scale processes that generate microbial diversity requires sampling strategies and analyses that enable very fine resolution (22, 23).

## Methods

### Genome assembly and annotation

*Pseudonocardia* strains were isolated from the cuticle of fungus-growing ants (24) and sequenced using either Pacific Biosciences technology at Duke University (EC080625-04) or Illumina at Washington University in St. Louis. PacBio assemblies were performed using HGAP 1.4 (25), while Illumina genomes were assembled using Velvet (26). Protein coding genes for all genomes were predicted de novo using Prodigal v2.60 (27), while ribosomal RNAs were predicted using RFAM (28) hidden Markov models and Infernal 1.1.1 (29). The origin of replication was predicted using OriLoc (30). Protein coding genes were annotated using TIGRFam v15 (31), PFAM v29, KEGG (32, 33), and actNOG (34) hidden Markov models via HMMer 3.1 (35). Natural product biosynthesis clusters were identified in each genome by antiSMASHv3 (36) followed by manual curation of cluster boundaries. Natural product gene clusters were grouped into families by 80% nucleotide identity via nucmer (37) alignment and 50% coverage for each segment of a cluster that matched another. Core and pan genome analyses were conducted using actNOG gene family annotation. All pathway and functional enrichment analyses were based on KEGG annotations.

### Population clustering

SNP identification was performed by mapping all genome assemblies to the complete genome of EC080625-04 using nucmer. Reference SNP positions that were covered by a contig for every genome were used for clustering using fineSTRUCTURE (38). Multiple runs with varying values of c and estimated population size had little effect on overall strain clustering, except for very high values of c which caused neighboring clusters to merge (i.e. merging clusters BCI-A and BCI-B into a single cluster).

### Recombination analysis

Recombination events were inferred using bratNextGen (39) on the nucmer alignments of all genomes to EC080625-04, for 10 iterations and 100 replicates with a significance cutoff of 0.01. The alpha value was reported as 2.7326. Statistically significant deviations from a random distribution of recombination events were found by comparing the number of recombination events that affected a window to a binomial distribution. A random distribution of recombination would result in each window being affected by approximately 4 events. Blocks of the chromosome that displayed high recombination density were defined as sets of 30 consecutive windows of 500bp each where the median fold enrichment of recombination events was at least 4 fold.

### Conserved gene analysis

Tajima’s D and monophyletic gene analyses were conducted using genes from EC080625-04 and the matching regions in contigs from other genomes aligned to this reference. Tajima’s D was calculated for genes that had at least 1% polymorphic nucleotide sites within each population. Monophyletic genes were identified by generating nucleotide alignments for all genes in EC-080625-04 using MAFFT v7.221 (40), followed by gene phylogenies generated using FastTree 2.0 (41). KEGG category enrichment for each lineage was determined using Fisher’s Exact Test and a Benjamini-Hochberg false discovery rate of 10%.

### Data Accessibility

Genome sequences and annotations are available at DOI: 10.5281/zenodo.2560835

## Acknowledgments

This project was supported through National Institutes of Health (NIH) U19 Al109673, NIH U19 TW009872, CAREER Award DEB-747002, National Science Foundation (NSF) MCB-0702025, and NSF MCB-0731822. Additional support was provided to MGC through NIH National Research Service Award T32 GM008505.

---

**Supplemental Figure1:**
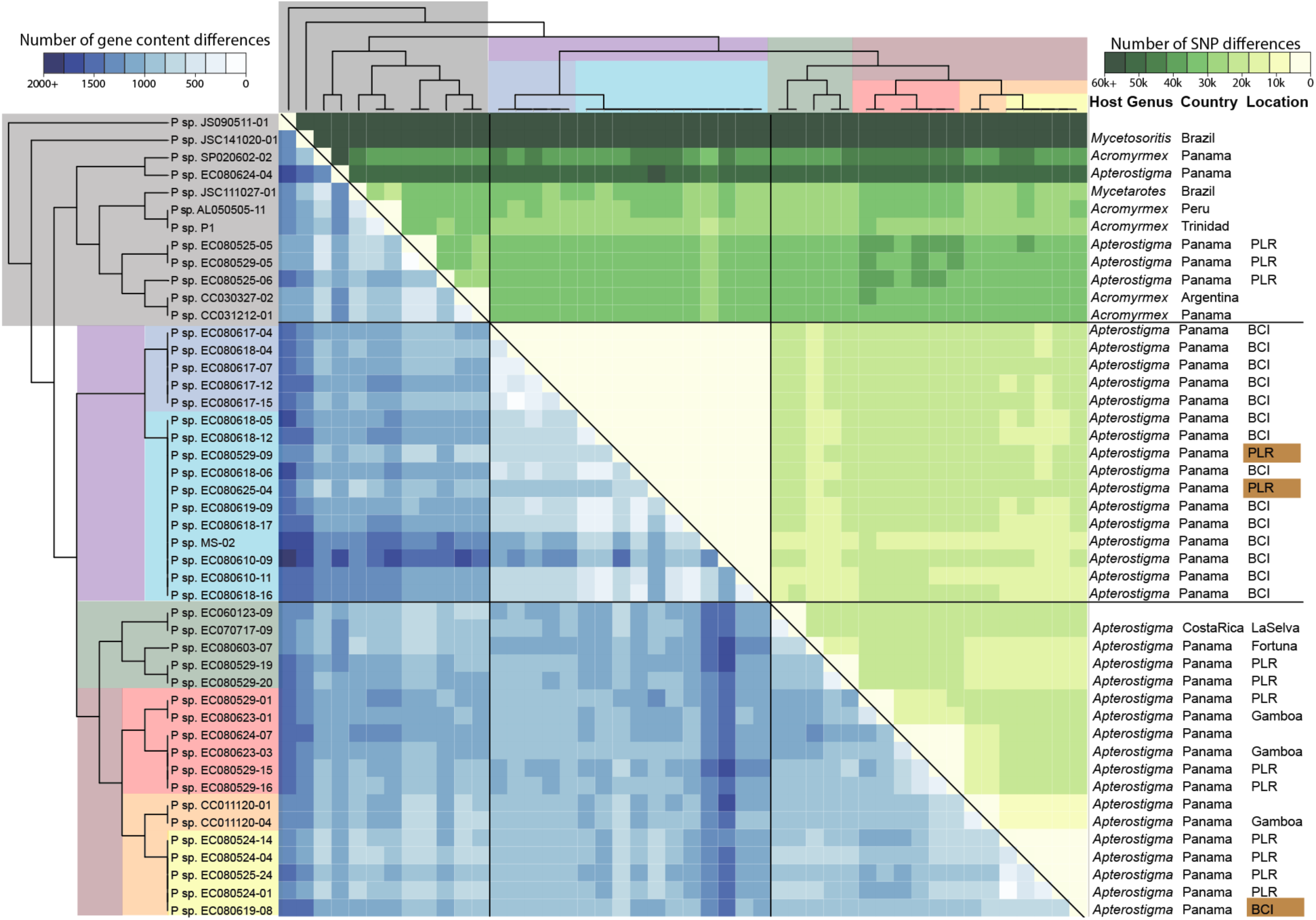
Genome content and SNP diversity in ant-associated *Pseudonocardia*. The number of actNOG gene content differences (blue) or SNP differences (green) between genome pairs. Isolation hosts and locations are given on the right.

**Supplemental Figure2:**
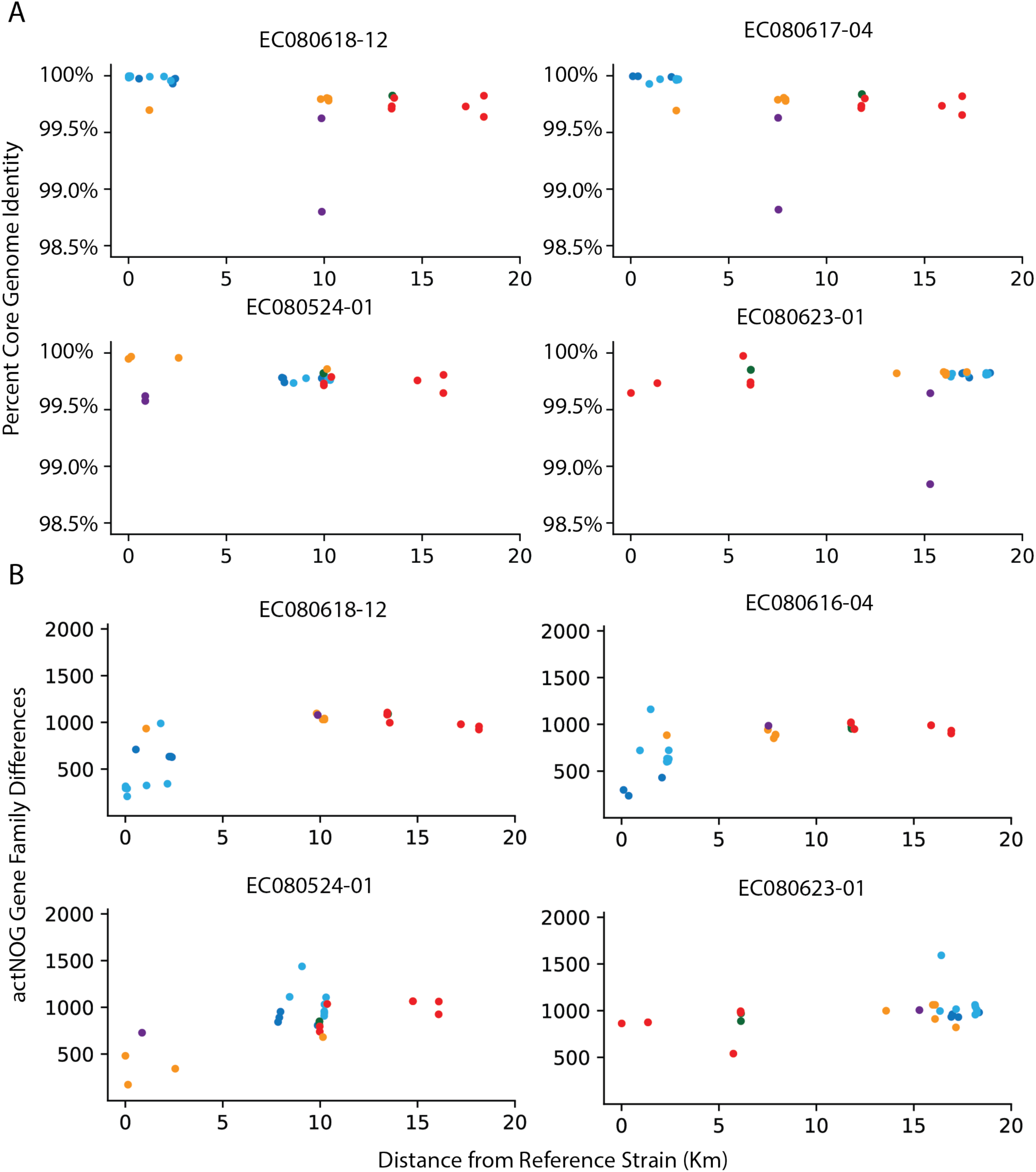
Conserved gene divergence and gene content divergence across geographic space. A. Core gene percent identity between PacBio assembled reference strains and all strains with GPS coordinates. Points are colored by strain population. Plot titles indicate the strain to which all other strains are compared. B. Raw number of gene content differences between PacBio finished reference strains and all others with GPS coordinates.

**Supplemental Figure3:**
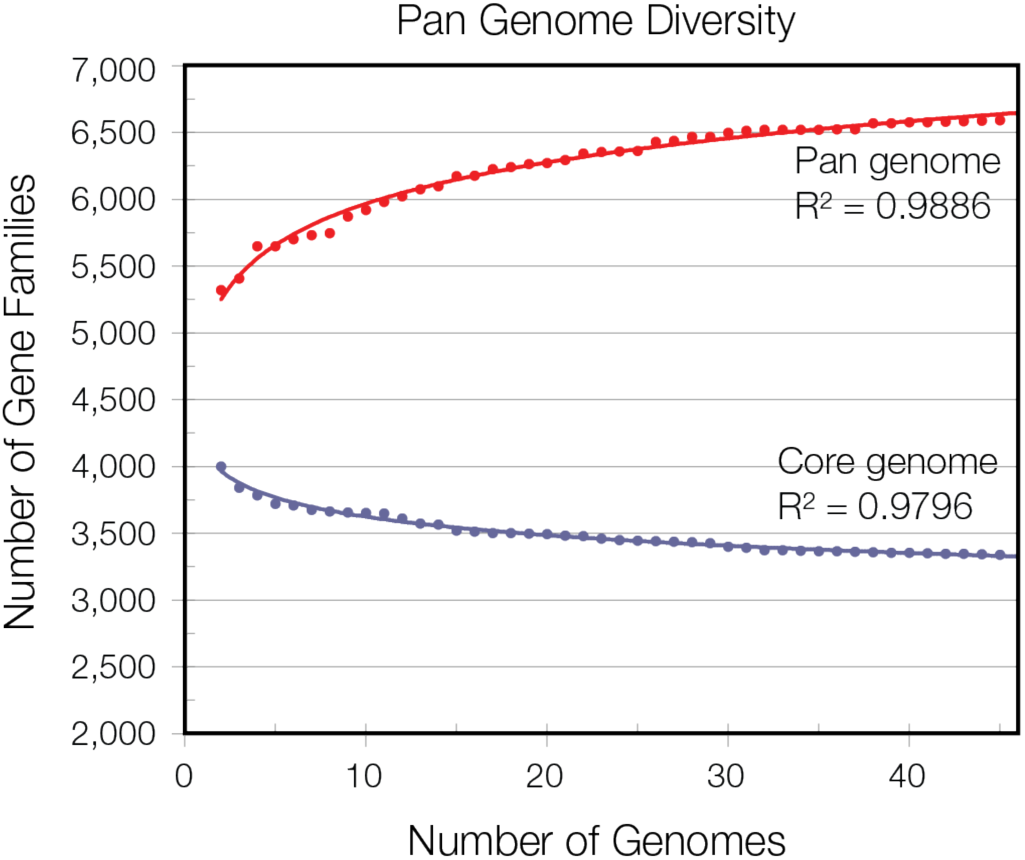
Pan-genome and core-genome size in closely related ant-associated *Pseudonocardia*. The total number of actNOG gene families present in the genome dataset (red) and the number of actNOG gene families conserved in all genomes (blue), as more genomes are added to the dataset. The pan-genome size fits a logarithmic function, while the core-genome size fits a power-law function.

**Supplementary Table 1.** Genome assembly and annotation statistics

| Genome | GenomeSize | N50 | TotalContigs | GC | GeneCount |
| --- | --- | --- | --- | --- | --- |
| AL050505-11 | 6566921 | 6389648 | 3 | 73.78 | 6046 |
| CC011120-01 | 6369072 | 5458 | 2178 | 73.55 | 7396 |
| CC011120-04 | 6368325 | 6216 | 1884 | 73.56 | 7258 |
| CC030327-02 | 5921325 | 6235 | 1740 | 73.72 | 6891 |
| CC031212-01 | 5889746 | 7200 | 1736 | 73.41 | 6824 |
| EC060123-09 | 6201792 | 8217 | 1463 | 73.74 | 6881 |
| EC070717-09 | 6351919 | 10828 | 1128 | 73.79 | 6753 |
| EC080524-01 | 6189181 | 9548 | 1282 | 73.61 | 6633 |
| EC080524-04 | 6371034 | 13375 | 903 | 73.8 | 6459 |
| EC080524-14 | 6195412 | 10360 | 1087 | 73.73 | 6485 |
| EC080525-05 | 6068818 | 9480 | 1349 | 73.88 | 6586 |
| EC080525-06 | 6836013 | 9998 | 1385 | 73.45 | 7328 |
| EC080525-24 | 6269987 | 6880 | 1629 | 73.54 | 7030 |
| EC080529-01 | 7106933 | 8171 | 1728 | 73.16 | 7892 |
| EC080529-05 | 6103212 | 8937 | 1390 | 73.85 | 6681 |
| EC080529-09 | 6348069 | 6770 | 1714 | 73.56 | 7171 |
| EC080529-15 | 6433317 | 10850 | 1259 | 73.64 | 6870 |
| EC080529-16 | 6427229 | 10977 | 1210 | 73.6 | 6783 |
| EC080529-19 | 6745963 | 8650 | 1626 | 73.4 | 7452 |
| EC080529-20 | 6861062 | 12349 | 1260 | 73.43 | 7207 |
| EC080603-07 | 6481475 | 11157 | 1109 | 73.78 | 6766 |
| EC080610-09 | 7131853 | 6138223 | 3 | 73.34 | 6597 |
| EC080610-11 | 6787135 | 4988 | 2501 | 72.94 | 8225 |
| EC080617-04 | 6932605 | 9808 | 1412 | 73.35 | 7393 |
| EC080617-07 | 6939821 | 11292 | 1271 | 73.29 | 7331 |
| EC080617-12 | 6879092 | 6338 | 2033 | 73.21 | 7935 |
| EC080617-15 | 6789668 | 8761 | 1475 | 73.31 | 7324 |
| EC080618-04 | 6789668 | 8761 | 1475 | 73.31 | 7324 |
| EC080618-05 | 7101947 | 12991 | 1127 | 73.21 | 7359 |
| EC080618-06 | 7107275 | 16463 | 990 | 73.23 | 7295 |
| EC080618-12 | 7021027 | 7448 | 1858 | 73.12 | 7878 |
| EC080618-16 | 6923749 | 5242 | 2468 | 72.99 | 8357 |
| EC080618-17 | 7014064 | 7082 | 1842 | 73.15 | 7863 |
| EC080619-08 | 6766081 | 5881 | 2090 | 73.27 | 7783 |
| EC080619-09 | 7024805 | 14182 | 1039 | 73.28 | 7249 |
| EC080623-01 | 6785441 | 6527 | 1963 | 73.3 | 7798 |
| EC080623-03 | 6881261 | 10164 | 1334 | 73.35 | 7321 |
| EC080624-04 | 6961708 | 7831 | 1962 | 73.21 | 7795 |
| EC080624-07 | 6855742 | 6063 | 2110 | 73.26 | 7919 |
| EC080625-04 | 6135769 | Na | 1 | 73.84 | 5744 |
| JS090511-01 | 6608484 | 5335 | 2398 | 73.85 | 8033 |
| JSC111027-01 | 6362052 | 6342421 | 2 | 73.95 | 5825 |
| JSC141020-01 | 7303443 | 6658632 | 4 | 73.62 | 6619 |
| MS-02 | 6188306 | 3359 | 3092 | 72.33 | 8189 |
| P1 | 6388771 | 14149 | 875 | 73.25 | 6636 |
| SP020602-02 | 6322523 | Na | 1 | 74 | 5771 |
| MedianAll | 6677223 | 8849 | 1401 | 73.42 | 7253 |
| MedianHighQuality | 6711593 | 6366034 | 2.5 | 73.81 | 5935 |

